# Explaining attractive and repulsive biases in the subjective visual vertical

**DOI:** 10.1101/2025.09.18.677071

**Authors:** Stefan Glasauer, W. Pieter Medendorp

**Author notes:** Correspondence: Stefan Glasauer.

## Abstract

Perception of gravity can be assessed by measuring the subjective visual vertical (SVV), the visually indicated spatial direction that appears earth-vertical to an observer. When the SVV is measured in darkness while the observer is roll-tilted, it shows substantial biases. At tilts larger than 45°, the bias is attractive, that is, the visual indicator appears vertical when rotated toward the observer. At smaller tilts, however, a repulsive bias is observed. The attractive bias has been explained within the Bayesian framework as the effect of a prior for upright posture. The repulsive bias has so far been considered anti-Bayesian, suboptimal, or as the result of uncompensated ocular counterroll. Here we show that both biases can be explained within a purely Bayesian model. More specifically, the repulsive bias at small roll-tilts is a consequence of the known tilt-dependent variability of the SVV, which is hypothesized to reflect different levels of sensory noise of the otolith organs. We thus provide a solution to a century-old question of why there is a repulsive bias in vertical perception.

## Introduction

Systematic biases in perception can be either attractive or repulsive with respect to the immediate spatial or temporal context (e.g., Wei & Stocker 2015, Fritsche et al. 2020, Akselberg et al. 2025). In some cases, both types of biases exist in combination. Such a case occurs for the visual perception of the direction of gravity, the so-called subjective visual vertical (SVV). The SVV is experimentally assessed by the participant adjusting a visual line in front of her to the perceived direction of gravity while being tilted in various roll-tilt positions (Fig. 1). From the very beginning of studying the SVV, researchers have noted that in the upright position, the SVV error is close to zero (Fig. 1A), while with large tilt angles around 90°, the SVV tilts toward the participants own head axis and thus lies between the true gravitational vertical and the head-centered vertical (Fig. 1C). In other words, it is attracted toward the head axis. However, for small tilt angles and some of the participants, the SVV is adjusted on the opposite side of the gravitational vertical, that is, it is repulsed from the head axis (Fig. 1B). The attractive bias has been named Aubert or A-effect (Aubert 1861), the repulsive bias is referred to as Müller or E-effect (Müller 1916), each after the researchers first reporting them. Even though the Müller effect has been found in many studies since (e.g., Van Beuzekom & Van Gisbergen 2000) and was even demonstrated in non-human primates (Daddaoua et al. 2008), the reason for this over-estimation of tilt is yet unclear. This is the focus of the present paper.

**Figure 1:**
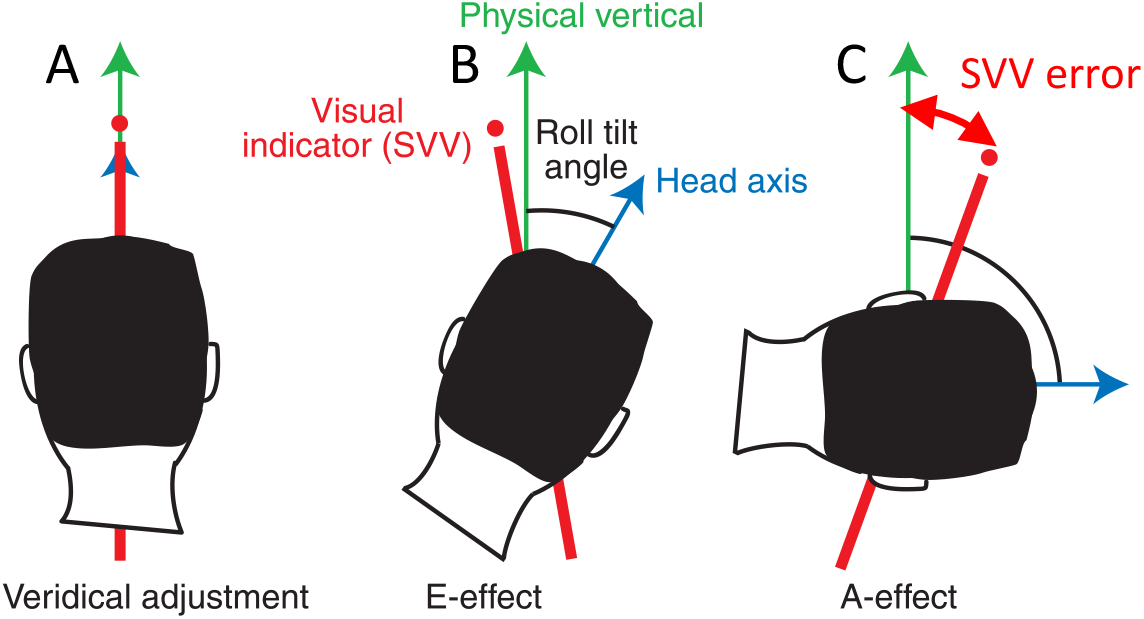
Schematic representations of biases in the SVV task (modified from De Vrijer et al. 2009). **A**: no bias while upright. **B**: Small body tilt: repulsive bias (E-effect). **C**: Larger body tilt: attractive bias (A-effect). Error: angle between physical and subjective vertical direction.

One of the first attempts of modelling the adjustment of the SVV (Mittelstaedt 1983) was based on a vectorial hypothesis, where a head-fixed vector pointing in the direction of the head axis, called the idiotropic vector, was added to the the gravity vector as measured by the otolith organs in the inner ears. The idiotropic vector alone yielded a good fit of the attractive A-effect but could not account for the E-effect. Therefore, Mittelstaedt (1983) hypothesized that the relative weight of the two otolith organs, the utricles and the saccules, was not equal but larger for the utricles. Together with a normalization this hypothesis could account quite well for both the E-effect and the A-effect (Fig. 2). Although this model was highly successful in quantitatively fitting the SVV, it does not offer a normative explanation of why perceptual biases of the SVV occur.

**Figure 2:**
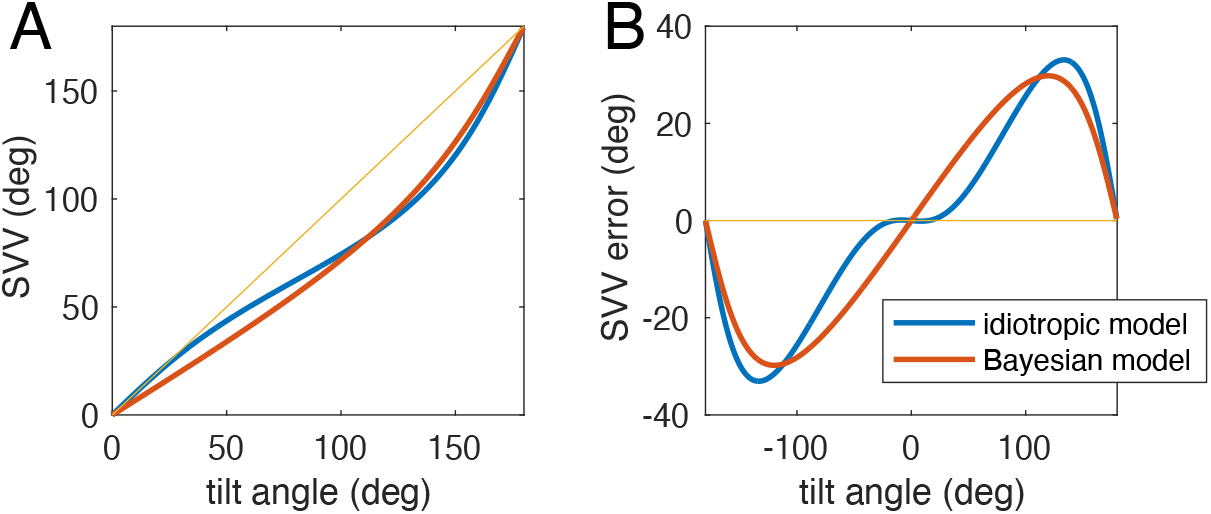
The vectorial idiotropic model (blue) after Mittelstaedt (1983) and a Bayesian model with a head-fixed prior for upright position. The parameters of the Bayesian model (see Methods) have been adjusted to fit the idiotropic model. A: SVV plotted over tilt angle for roll tilt between upright (0°) and upside down (180°). B: SVV error plotted over tilt angle for the full circle. The repulsive E-effect simulated by the idiotropic model is clearly visible.

With the advent of Bayesian inference approaches and conceiving perception as a close-to-optimal estimation process, the view on apparent perceptual errors and biases in the SVV also changed. The question was posed whether errors in the SVV are the result of a normative Bayesian estimation process, which aims to optimize the estimation in the presence of uncertainty such as sensory measurement noise. Within the Bayesian perspective, an obvious reason for an attractive bias such as the A-effect is that head-upright is the most common orientation with respect to gravity, which can be modelled as a prior distribution centered at roll-tilt zero, i.e., upright. Perceptual estimates would then be attracted towards this prior, resulting in the A-effect (see Fig 2). The first Bayesian model for the SVV was proposed by a student of Mittelstaedt (Eggert 1998), who successfully could explain the A-effect as consequence of a head-fixed prior for upright position. The Bayesian approach has since then been used by many authors to explain various features of the perception of gravity (Van Beuzekom & Van Gisbergen 2000, MacNeilage et al. 2007, De Vrijer et al. 2008, De Vrijer et al. 2009, Clemens et al. 2011, Alberts et al. 2016, Willemsen et al. 2022, and many more). Despite this success, one problem has remained: explaining the repulsive bias of the E-effect with a Bayesian framework. The usual Bayesian approach predicts that estimated values are attracted towards the prior but not repelled by it. Eggert’s model, for instance, attributed the E-effect to suboptimal integration of differences in measurement noise of the two otolith organs, without providing a complete Bayesian account. Other studies have linked the E-effect to an uncompensated effect of ocular counterroll (e.g., De Vrijer et al. 2008) or neck proprioception (Clemens et al. 2011) with the visual adjustment of the SVV, yet others have neglected it. Fig. 2 shows a comparison between the original idiotropic model and a Bayesian model with a head-fixed prior in order to demonstrate that the A-effect, but not the E-effect, can be simulated. Consequently, repulsive biases such as the E-effect have also been called “anti-Bayesian” in the literature (e.g., Brayanov & Smith 2010, Wei & Stocker 2015, Peters et al. 2016).

There have been different approaches to explain anti-Bayesian or repulsive biases in literature (e.g., Schwartz et al. 2007, Girshick et al. 2011, Wei & Stocker 2015) using a variety of mechanisms, ranging from sensory adaptation over efficient coding to various flavors of probabilistic estimation, variably called Bayesian or predictive or optimal. However, none of these approaches have yet been applied and tested to explain the E-effect in the perception of the SVV.

In the following, we show that the E-effect can be explained as Bayes-optimal and in accordance with Bayesian perception, if, as proposed previously (Eggert 1998), sensory noise for both otolith organs is assumed to differ. We also show that such a model is in accordance with the well-known anisotropy of the variability of the SVV (e.g., Lechner-Steinleitner 1978, Van Beuzekom & Van Gisbergen 2000, Tarnutzer et al. 2009).

## Methods

Participants in typical experiments are asked to adjust a visual indicator, usually a line, in an otherwise dark environment to the direction which they perceive as upright. This adjustment is performed in different roll-tilt body positions (see Fig. 1). We therefore implemented a Bayesian model of the SVV, for which we considered as input the vestibular information from the otolith organs, which measures the direction of gravity, and as output the perceived tilt of the gravitational upright, which is provided as roll position of the visual indicator.

### The Bayesian model

For the Bayesian model we consider that the true roll-tilt angle of the head with respect to gravity (Fig. 1) is unknown but can be measured by the vestibular sensors. However, this sensory measurement is corrupted by measurement noise. Therefore, the perceptual process aims to estimate the roll-tilt angle by combining prior knowledge about the distribution of head positions with knowledge of the measurement uncertainty. We assume that the prior knowledge for the angular head tilt can be represented as von-Mises distribution with mean 0 and dispersion 1/*κ*_*h*_. The measurement uncertainty comes from a von-Mises distribution with mean 0 and dispersion 1/*κ*_*v*_. Importantly, we assume that the measurement noise can be signal-dependent, that is, the dispersion 1/*κ*_*v*_ depends on the actual tilt angle.

The Bayesian estimation model for this case is formalized as follows. The prior distribution is given as von-Mises probability density function:

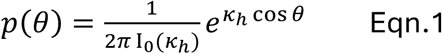

with *θ* being the roll-tilt angle of the head and I_0_(*κ*_*h*_) the modified Bessel function of the first kind of order 0 (required for normalizing the probability density).

The measurement distribution, i.e., the distribution of the measurement *φ* for a given stimulus value *θ* is:

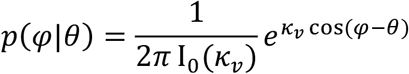

The likelihood function is the measurement distribution viewed as function of the parameter (Ma et al. 2024), the unknown roll-tilt angle of the head *θ*:

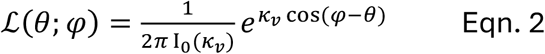

with *φ* being the measured angle.

The posterior distribution can now be calculated by multiplying the prior distribution and the likelihood function, and normalizing it so that the integral over the posterior density becomes unity:

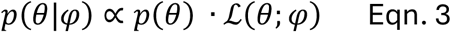

Finally, as estimated or perceived tilt angle, the SVV, the angular expectation of the posterior density *p*(*θ*|*φ*) is computed.

#### Bayesian model 1: constant sensory noise

If the sensory noise is independent of roll tilt angle and thus *κ*_*v*_ is constant, then the likelihood is a von-Mises probability density, and the resulting posterior distribution is again a von-Mises density. It is possible to analytically provide the posterior (not shown, see for example Tarnutzer et al. 2009). The two free parameters of this constant noise model are *κ*_*h*_ and *κ*_*v*_.

#### Bayesian model 2: signal-dependent noise, full likelihood

For signal-dependent noise, the dispersion 1/*κ*_*v*_ depends on the true tilt angle *θ*, so that the likelihood function becomes

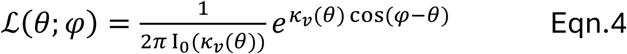

Note that in general this is no longer a von-Mises density. However, it becomes evident that an accurate likelihood function now requires complete knowledge of the noise characteristics at any possible tilt angle.

To implement signal-dependent noise, we consider the measurement of the tilt angle by the vestibular gravity sensors, utricles and saccules. The utricles measure the sine, the saccules the cosine of the roll tilt angle *θ* with respect to gravity. We assume that both sensors are affected by constant measurement noise ε_*x*_ and ε_*z*_ which is normally distributed with zero mean and variances 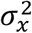 and 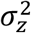. We allow the noise variances of utricles and saccules to be different as proposed previously (Eggert 1998). The measured angle *φ* is thus

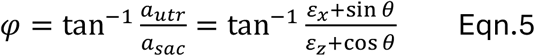

with measured accelerations *a*_*utr*_ and *a*_*sac*_ given in units of *g* with *g* = 9.81 *m/s*^2^. Using a first-order Taylor series approximation for error propagation, the expectation of the measured angle *φ* equals the actual roll angle *θ*, and we can approximate the resulting variance of the angle *φ*, and therefore the noise dispersion 1/*κ*_*v*_(*θ*), as

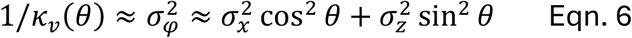

Thus, if 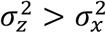, the noise dispersion is minimal for 0° (upright) and 180° (upside down), with maxima for 90° and −90° roll tilt. The signal-dependent noise model has three free parameters, the noise variances 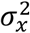 and 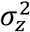, and the concentration parameter of the prior *κ*_*h*_.

#### Bayesian model 3: signal-dependent noise, local likelihood approximation

Instead, we can also choose an approximative strategy by assuming that noise dispersion depends on the measured angle *φ* rather than the true tilt angle *θ*. In that case, only the local information about noise dispersion at the particular measured tilt angle is required. The resulting approximative likelihood function is again a von-Mises density, even though a different one for each measurement:

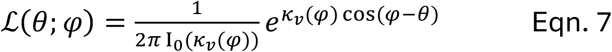

with 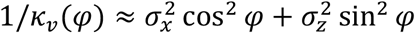 and thus again three free model parameters.

### The idiotropic model

We used the idiotropic model of the SVV (Mittelstaedt 1983) as comparison, because it provides a good description of the known data and can thus be used as reference. It is given by the following equations. For consistency with Mittelstaedt’s original publication, the SVV angle is denoted as *β*. The tilt angle *θ* is measured as otolith responses of the utricles *u* and saccules *s*.

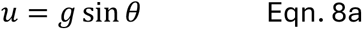

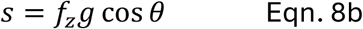

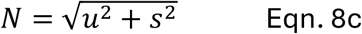

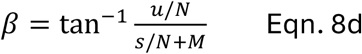

where *g* = 9.81 m/s^2^ is gravitational acceleration, *f*_*z*_ the saccular weighting, and *M* the idiotropic vector. *N* is a normalization factor. Mittelstaedt’s original parametrization was *M* = 0.4 and *f*_*z*_ = 0.7, producing a large A-effect. For the simulation of the idiotropic model in Fig. 2 and demonstration of the E-effect, we these original parameters.

### Response variability

The variability of the posterior distribution is, in general, not equal to the variability of the distribution of behavioral responses (Clemens et al. 2011; Ma et al. 2022). While the variability of the response distribution can be analytically derived in the Gaussian case, this is not possible for the non-linear function relating tilt angle to SVV. However, there are two possible solutions: since the SVV dependence on tilt angle in the models is a deterministic function,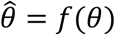 the SVV response variance can be approximated via 1^st^ order Taylor expansion by 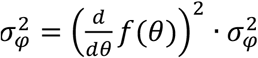, with 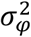 being the variance of the measurement noise (Eqn. 6). Alternatively, the variance of the SVV at a particular stimulus angle can be estimated by Monte-Carlo simulation. For the response variance shown in the Results, we chose the latter and simulated the model n=1e5 times for each tilt angle with noisy measurements according to Eqn. 5.

### Model simulation and fitting

Models were simulated numerically using Matlab (The Mathworks) according to the equations above with a discretization step of 2*π*/1000 for representing densities and likelihoods. For fitting the Bayesian models to the idiotropic model and to the data, we used the Matlab function lsqnonlin. Differences between two angles *φ*_1_ and *φ*_2_, required for fitting, were calculated using complex numbers as arg 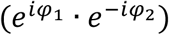. As mentioned above, the SVV angle was computed as angular expectation of the posterior density of the respective models. The standard deviations of the posterior and the response distributions were calculated as square root of the circular variance.

## Results

We first simulated the idiotropic model with original parameters *M* = 0.4 and *f*_*z*_ = 0.7 (Mittelstaedt 1983) as reference. We then fitted the SVV angles predicted by the three Bayesian models to the SVV angles of the idiotropic model. As shown in Fig. 3A and B, the model with signal-dependent noise and full likelihood can simulate an E-effect and closely resembles the idiotropic model, unlike the other two Bayesian models. As expected from visual inspection, the root-mean-square error for the full likelihood model was much smaller (0.6°) than for the models with local likelihood approximation (5.3°) or with constant noise (6.0°).

**Figure 3:**
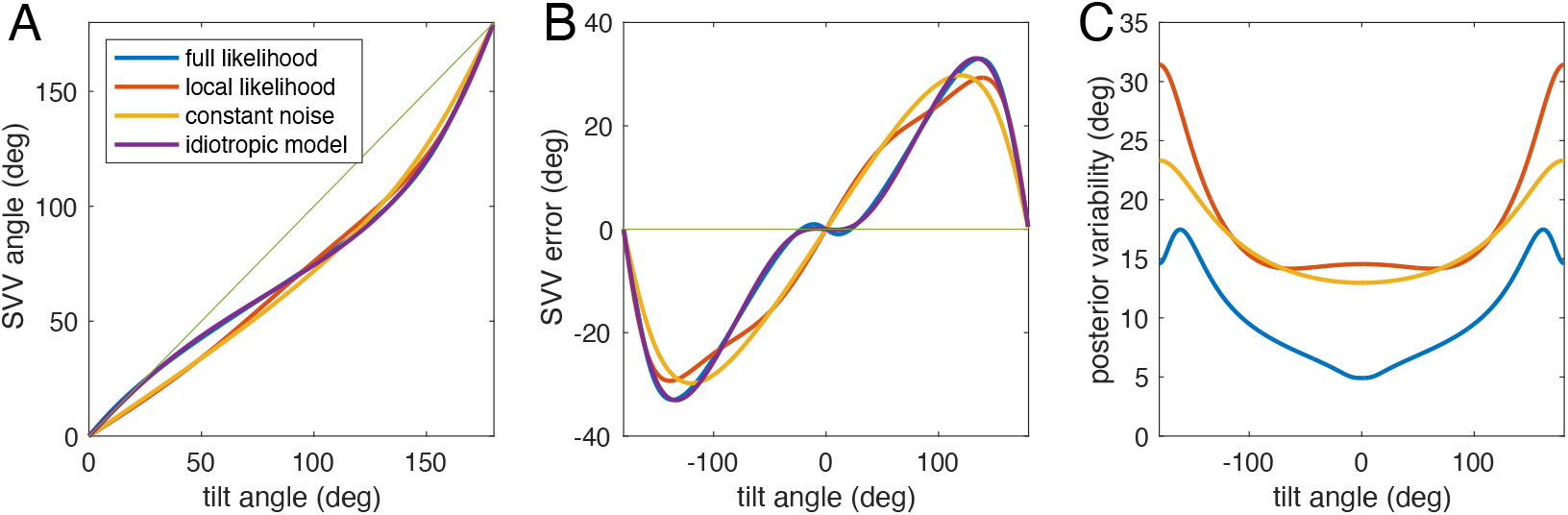
Comparison of the idiotropic model (violet) after Mittelstaedt (1983) and the best fits of the three Bayesian models to the idiotropic model. A: SVV angle plotted over tilt angle for roll tilt between upright (0°) and upside down (180°). B: SVV error plotted over tilt angle for the full circle. The E-effect is visible for the idiotropic model and the Bayesian model with full likelihood, but not for the other two Bayesian models. C: predicted posterior variability (square root of circular variance) for all three Bayesian models. Without additional assumptions, the idiotropic model does not offer a prediction for variability. Note that the actual variability of SVV responses can differ from the posterior variability (Clemens et al. 2011, Ma et al. 2022).

Fig. 3C shows the predicted standard deviation of the SVV for the three Bayesian models. The predictions have in common that the standard deviation should be small at the upright position and larger at tilted positions, which is indeed the case (e.g., Van Beuzekom & Van Gisbergen 2000, Tarnutzer et al. 2009). The model parameters of the full likelihood model were 1/*κ*_*h*_ = 0.17 (corresponding to an angular standard deviation of the prior 24.8°), utricular standard deviation σ_*x*_ = 0.09 g, and saccular SD σ_*z*_ = 0.33 g. To visualize the complete estimation process, inspired by Girshick et al. (2011), Fig 4 depicts our full likelihood model, using the parameters given above.

**Figure 4:**
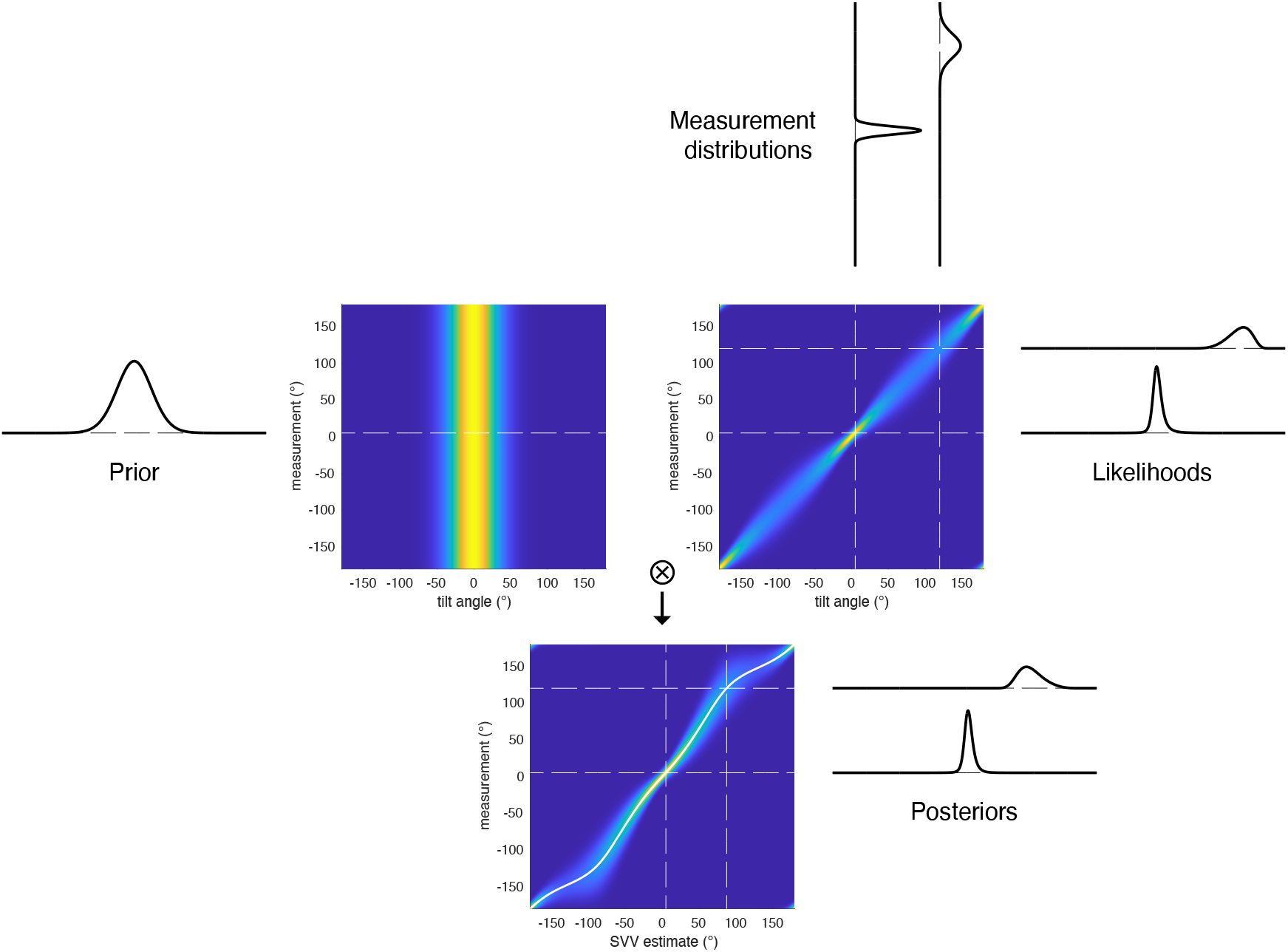
Schematic for the Bayesian estimation process of the full likelihood model (adapted to the current model after Girshick et al. 2011). In the colored panels, the x-axis is physical tilt angle (upper panels) or estimated SVV angle (lower panel) and the color corresponds to probability. The left upper panel show the prior, which is independent of the measurement. Vertical slices through the upper right panel show the symmetric measurement distributions, horizontal slices are the likelihood functions, which become asymmetric. Element-wise multiplications of these matrices (⊗) and row-wise normalization of the result yields the lower panel, which gives the posterior distributions as horizontal slices. The angular mean of the posterior distributions is plotted as white line in the lower panel. Note that axes in this lower panel are flipped in comparison to Fig. 3A. Slices are shown for tilt angles of 5° and 120°, parameters are computed from fitting the idiotropic model.

For a comparison with real data, we digitized Figs. 4 and 11 of Van Beuzekom & Gisbergen (2000), who measured the SVV and its variability in six subjects. We then fitted the full-likelihood model to the SVV data and predicted the variability. Both results are shown in Fig. 5. As can be seen from the figure, the fit captured the SVV data very well, and the *predicted* variability is not only in the same range as the SVV variability reported by Van Beuzekom & Gisbergen (2000), but also approximately matches the shape of the variability dependence on roll tilt. The model parameters for this fit are the dispersion of the prior 1/*κ*_*h*_ = 0.24 (corresponding to a circular standard deviation of 30.6°), the utricular standard deviation σ_*x*_ = 0.07 *g*, and the saccular SD σ_*z*_ = 0.34 *g*.

**Figure 5:**
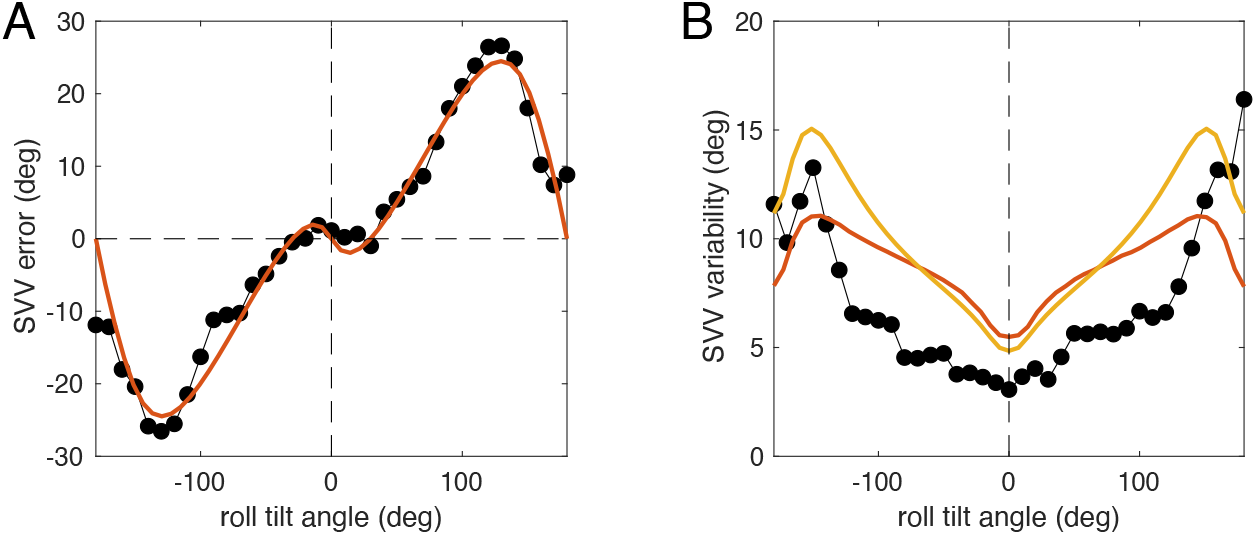
The full-likelihood model (red) and data (black dots) from Van Beuzekom & Gisbergen (2000). A: SVV error (black) plotted over roll tilt angle together with best fit of the model (red). B: SVV variability (black) plotted over roll tilt angle together with predicted response variability (red) and variability of the posterior distribution (yellow) from numerical model simulation, both expressed as square root of the circular variance.

To understand what determines whether an E-effect occurs or not, the model comparisons in Fig. 3 are helpful. The E-effect could be fitted only with the full-likelihood model with signal-dependent noise, but not if the likelihood functions were approximated by the local noise distributions. In Fig. 6 we compare noise distributions with likelihood functions: while noise distributions are symmetric, likelihood functions for small tilt angles are asymmetric (see also Fig. 4).

**Figure 6:**
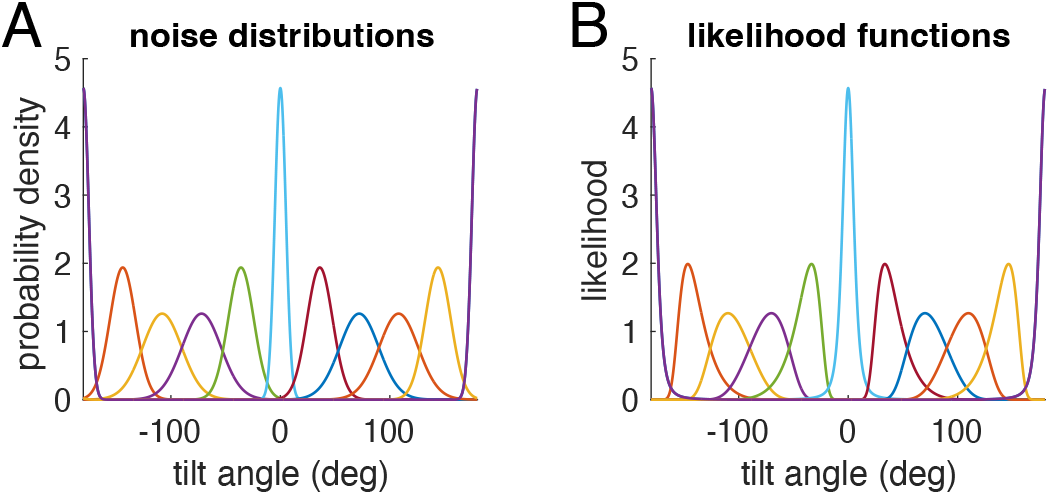
Noise distributions (A) and corresponding likelihood functions (B) for the full-likelihood model shown in in Fig. 3 and 4. Each color shows one noise distribution centered at the tilt angle in A and the corresponding likelihood function in B. While the noise distributions are symmetric von-Mises distributions, the likelihood functions for small tilt angles are clearly asymmetric.

For modelling the perceived SVV angle as result of a Bayesian estimation, the likelihood function is multiplied with the prior distribution (Eqn. 3, and the mean of the resulting posterior distribution is considered the estimated SVV angle, see also Fig. 4). For both the local-likelihood model and the full-likelihood model, Fig. 7 shows all three distributions for a tilt angle of 10°. As evident from the figure, the resulting posterior distributions (yellow) are slightly different, and, more importantly, the mean of the posterior (dashed black line) is attracted towards zero for the local likelihood model (Fig. 7A) and thus smaller than the actual roll tilt, but it is repulsed from zero and larger than the tilt angle for the full-likelihood model (Fig. 7B).

**Figure 7:**
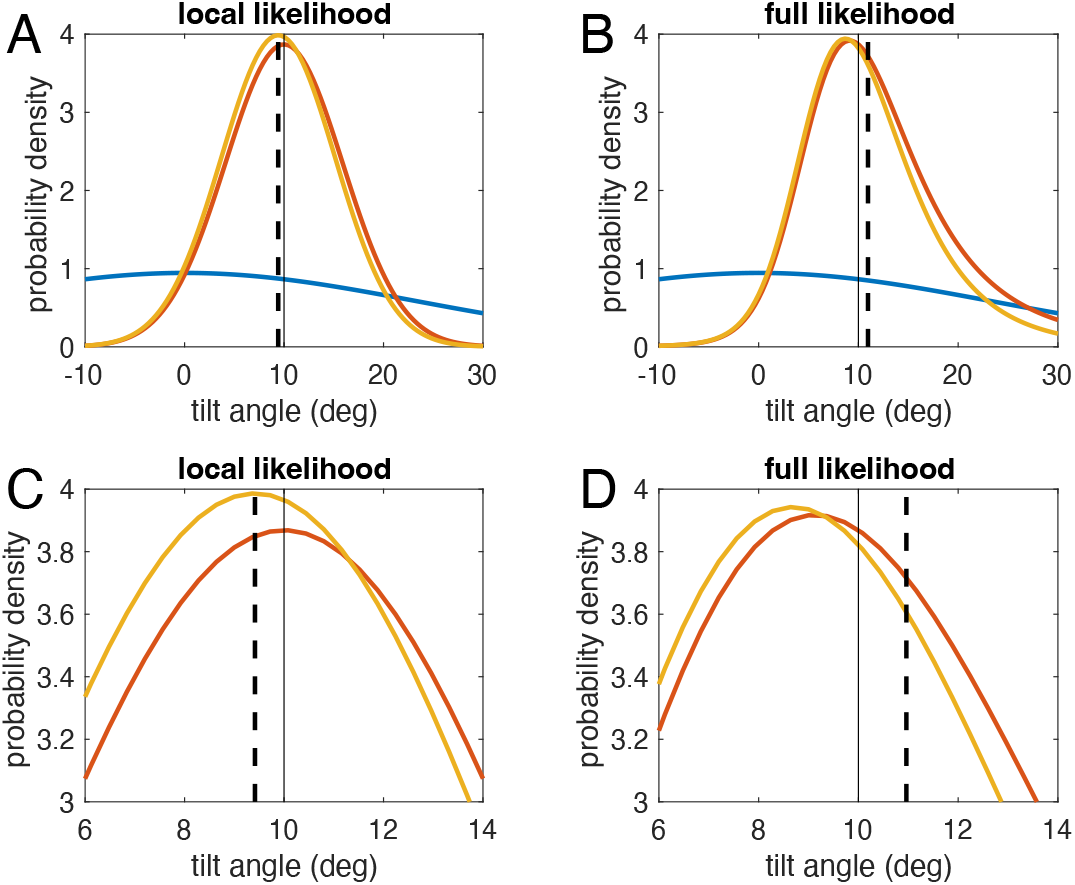
Bayesian estimation for two models: to estimate tilt angle, the measured angle (thin black line at 10°) is represented as likelihood function (red), which is multiplied with the prior density (blue). This yields the posterior density (yellow). The estimate is the mean of the posterior distribution (black dashed line). & and D are magnifications of A and B. A and C: For the local-likelihood model, prior, likelihood, and posterior are all symmetric. Thus, the posterior mean is attracted towards the prior mean (at zero). B and D: For the full-likelihood model, the likelihood function becomes asymmetric, leading to an asymmetric posterior distribution. Due to this asymmetry, the mean of the posterior is now larger than measured angle, which results in a repulsive effect. Note that only the full-likelihood model is optimal for the chosen signal-dependent noise distributions.

The ability of the model to simulate the E-effect thus depends on the asymmetry of the likelihood functions, which in turn depends on how much the measurement noise depends on the stimulus, i.e., the roll tilt angle. To visualize this dependence, we changed parameter 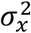 of the model (see Eqn. 6), the variance of the utricular measurement noise, from its fitted value to the same value as the saccular noise variance 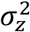. Figure 8 shows simulations of the resulting models. As evident from the SVV error, the model exhibits an E-effect only for strong stimulus dependence of the measurement noise. From the figure it can also be seen that the model can simulate SVV error with or without E-effect, depending on the relation between 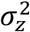 and 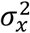.

**Figure 8:**
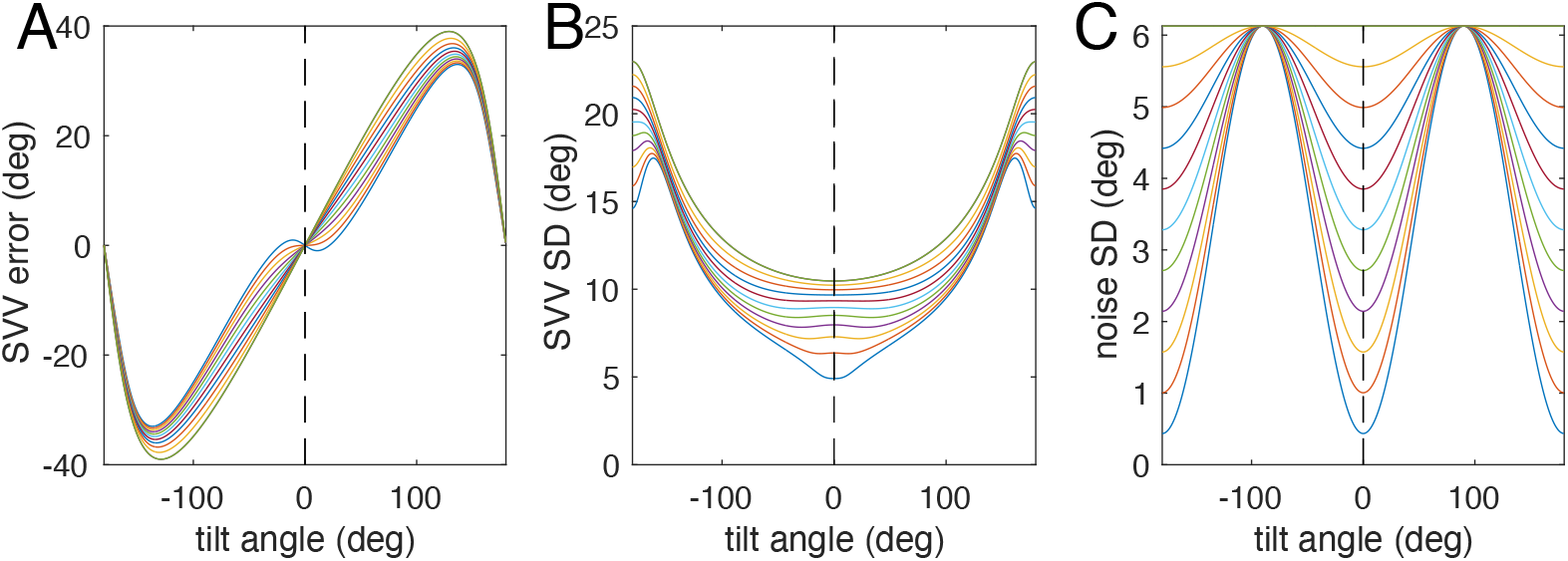
Dependence on the relation between the saccular noise variance 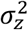 and the utricular noise variance 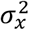 for the SVV error (A), the standard deviation of the posterior distribution, and the standard deviation of the measurement noise (C). The lowest thin blue line in B and & corresponds to the full-likelihood model shown in Fig. 3, 5, and 6B. The thin green line corresponds to equal variances and results in measurement noise independent of tilt angle dependence (top line in C) and no E-effect in A.

## Discussion

Here we show that the “anti-Bayesian” E-effect can be explained as Bayesian consequence of signal-dependent measurement noise resulting in asymmetric likelihood functions, which in turn causes the E-effect. Asymmetric likelihood functions are, in turn, a consequence of stimulus-dependent measurement noise, and can lead to repulsive bias (Girshick et al. 2011). Similar anti-Bayesian effects in orientation perception have been explained by asymmetric likelihood functions (Wei & Stocker 2015). While Wei & Stocker (2015) hypothesized efficient coding to be the reason for stimulus-dependent sensory noise, we showed here that the simple assumption of constant noise on the two otolith organs and its propagation through the internal computation of orientation with respect to gravity already causes signal-dependent sensory noise, which in turn results in an asymmetric likelihood function causing the repulsive bias of the E-effect.

We furthermore show that our full Bayesian model, having only three free parameters, can fit not only average SVV adjustments covering the full 360° range from the literature (Fig. 4A), but also closely mimics the variability found experimentally for these adjustments. However, a comparison with other datasets (e.g., Tarnutzer et al. 2009) revealed that while our model could fit the data, it usually predicted a variability which was higher than observed experimentally.

The result of the estimation stage of any Bayesian models is not a single value, but a posterior distribution, from which, in case of simulating a perceptual experiment like setting the SVV, a particular value must be chosen as estimate. Here we chose the mean of the posterior distribution, which minimizes the sum of the quadratic errors on a linear scale and the sum of the cosine distances on the circular scale. In the literature often the maximum of the posterior, the maximum-a-posteriori (MAP) estimate, is chosen. However, the MAP corresponds to a cost function that is 0 when the MAP exactly matches the stimulus and 1 otherwise. Such a 0-1 cost function is not well-suited for continuous estimation problems (see Ma et al. 2022), for example, because it means any error would be weighted equally, while it is more natural to put larger cost on larger errors. From Fig. 6B it is obvious that taking the MAP as SVV angle would, in the illustrated case, not lead to an E-effect, while the taking the mean does.

A recent comparison of SVV models (Willemsen et al. 2022), which focused on the effect of non-Gaussian priors, also tested a model that contained asymmetric likelihood functions but favored a previously proposed one with a Gaussian prior (Clemens et al. 2011), which attributed the E-effect to an interaction of proprioception with the visual adjustment of the SVV, but assumed a symmetric likelihood as modelled in our suboptimal local-likelihood model. While it is completely reasonable to posit proprioceptive influences to affect the SVV (and might explain why some of the SVV datasets in the literature show lower variability than predicted by our model), our present work aims to emphasize that even without such additional inputs an optimal Bayesian SVV adjustment is completely compatible with both E- and A-effects.

In the present model, the main reason for an E-effect is the asymmetric likelihood function, which in turn is caused by stimulus dependent noise. To model stimulus dependent measurement noise, we chose a very simple assumption: additive Gaussian measurement noise on both otoliths, which is stimulus-independent, but differs for utricles and saccules, being larger in the latter. This choice is consistent with anatomical evidence: the utricle contains substantially more hair cells than the saccule (Rosenhall, 1972). Because the utricle is functionally most sensitive to tilts around 0°, while the saccule is most responsive near 90° (Jaeger et al., 2008), this asymmetry may explain the increase in otolith noise at larger tilt angles (Tarnutzer et al., 2010).

Udo de Haes (1970) found stimulus independent variability of ocular counterroll, which would support our hypothesis of constant additive noise, since counterroll is determined predominantly by the utricles. Furthermore, neural variability of otolith afferents stays almost constant, at least for a range of ±0.5 g (Schneider et al. 2015), corresponding to ±30° tilt. Concerning differences in measurement noise, a recent study (Kobel et al. 2024) showed that perceptual thresholds of linear motion in the earth-horizontal plane along the utricular (y-) axis are much smaller (0.65 cm/s) than those along the saccular (z-) axis (1.2 cm/s). While this difference qualitatively supports our assumption, it would be interesting to see whether interindividual SVV characteristics correlate with differences in perceptual thresholds of linear motion.

An interesting consequence of the dependence of the E-effect on the complete likelihood function is that the stimulus dependence of the measurement distribution across the entire 360° range must be known for the estimation process. While it is conceivable for head tilt or other measures of orientation that the noise characteristics can be learned due to the closed circular range, it is practically impossible for magnitudes on open scales such as duration or distance. However, measurement of magnitudes on open scales does exhibit stimulus-dependent uncertainty. For duration perception this is known as scalar variability, and corresponds to Weber’s law, which holds for most other magnitudes, including distances. Consequently, one possibility for magnitudes on open scales would be that the brain uses Weber’s law as approximation of the true stimulus dependence of magnitudes, which in turn allows a good approximation of the respective likelihood functions. Such an approximation can be achieved by representing open-scale magnitudes on a logarithmic scale, as proposed previously (e.g. Petzschner et al. 2015).

In summary, our current model offers a Bayesian account of both the Aubert and Müller biases in the subjective visual vertical, which is optimal, if sensory measurement noise depends on tilt angle. As possible explanation for tilt-dependent noise, we showed that assuming additive Gaussian noise for both otolith organs is sufficient, if noise levels differ for utricles and saccules. Thus, our model can account for the repulsive bias known as the E-effect without requiring additional contributions from ocular counterroll or proprioception, as originally suggested by Müller (1916).

## Acknowledgements

WPM is supported by the following grants: NWA-ORC-1292.19.298, NWO-SGW-406.21.GO.009, and Interreg NWE-RE:HOME. SG received funding from Deutsche Forschungsgemeinschaft (project number 532480418).

